# Motor learning without movement

**DOI:** 10.1101/2021.08.12.456140

**Authors:** Olivia A. Kim, Alexander D. Forrence, Samuel D. McDougle

**Author notes:** Correspondence to: Olivia Kim. **AUTHOR CONTRIBUTIONS** Conceptualization, OK, AF, & SM; Methodology, OK, AF, & SM; Software, AF & OK; Validation, OK & AF; Formal Analysis, OK & AF; Investigation, OK & AF; Resources, OK & SM; Data Curation, OK & AF; Supervision, SM; Project Administration, SM; Funding Acquisition, OK & SM; Writing – original draft, OK; Writing – review & editing, OK, AF, & SM.

## Abstract

Prediction errors guide many forms of learning, providing teaching signals that help us improve our performance. Implicit motor adaptation, for instance, is driven by sensory prediction errors (SPEs), which occur when the expected and observed consequences of a movement differ. Traditionally, SPE computation is thought to require movement execution. However, recent work suggesting that the brain generates and accounts for sensory predictions based on motor imagery or planning alone calls this assumption into question. Here, by measuring implicit adaptation during a visuomotor task, we tested whether motor planning and well-timed sensory feedback are sufficient for SPE computation. Human participants were cued to reach to a target and were, on a subset of trials, rapidly cued to withhold these movements. Errors displayed both on trials with and without movements induced single-trial implicit learning. Learning following trials without movements persisted even when movement trials had never been paired with errors, and when the direction of movement and sensory feedback trajectories were decoupled. These observations demonstrate that the brain can compute SPEs without generating overt movements, leading to the adaptation of planned movements even when they are not performed.

**SIGNIFICANCE STATEMENT:** We are always learning from our mistakes, because the brain is constantly generating predictions and monitoring the world for any surprises, which are also referred to as “prediction errors.” Whenever a prediction error occurs, the brain learns to update future predictions and be more accurate. Here, we demonstrate that the brain predicts the consequences of movements, computes prediction errors, and updates future movements, even if we subsequently decide to withhold the movement. Thus, the brain can learn to update movements that are not performed, representing a mechanism for learning based only on movement planning and sensory expectation. These findings also provide further support for the role of prediction in motor control.

**SIGNIFICANCE STATEMENT:** Our brains control aspects of our movement without our conscious awareness – allowing many of us to effortlessly pick up a glass of water or wave “hello.” Here, we demonstrate that this implicit motor system can learn to refine movements that we plan but ultimately decide not to perform. Participants planned to reach to a target, and they sometimes withheld these reaches. When reaches were withheld, an animation simulating a reach that missed the target played. Afterwards, participants reached opposite the direction of the mistake without awareness of this change in their movements, indicating that the implicit motor system had learned from the animated mistake. These findings indicate that movement is not strictly necessary for motor adaptation, and that we can learn to update our actions based only on movement planning and observation of related events in the world.

## INTRODUCTION

Prediction errors help to optimize behavior by driving learning processes that correct for our mistakes. Accordingly, their computation is thought to be a fundamental feature of the nervous system (1). Specific types of prediction errors are associated with dissociable learning processes, with sensory prediction errors (SPEs) serving as the teachers of the implicit motor system. SPEs are thought to trigger the adaptation and refinement of movements when the predicted and expected sensory outcomes of a movement differ (2–5). Traditional formulations assume that movement execution is critical for SPE computation (6, 7). However, current thinking posits that the forward model estimates the consequences of movements before the relevant sensory feedback reaches the brain, thereby overcoming intrinsic physiological delays in sensory signal conduction to the brain and allowing for the rapid motor control required by most vertebrates (8). Taking this principle to its logical conclusion indicates that motor execution should not be necessary for the generation of predictions by a forward model, because movements are synchronous with the sensory outcomes that must be predicted before we can plan the next stages of movement. In other words, the sensory consequences of intended movements ought to be predicted before those movements occur, and movement itself should not be necessary for this predictive process.

Recent work offers indirect support for the claim that the brain might predict the sensory consequences of movements before they can be performed, even when the agent does not have a clear intention to move (9–13). Considering that sensorimotor prediction should not in theory require movement, it may be that a prediction can be combined with an observation to support SPE computation without any actual motor execution. That is, SPEs should be effectively computed based upon only two events – the generation of a sensory prediction and the observation of sensory feedback (**Fig. 1a**).

**Figure 1.**
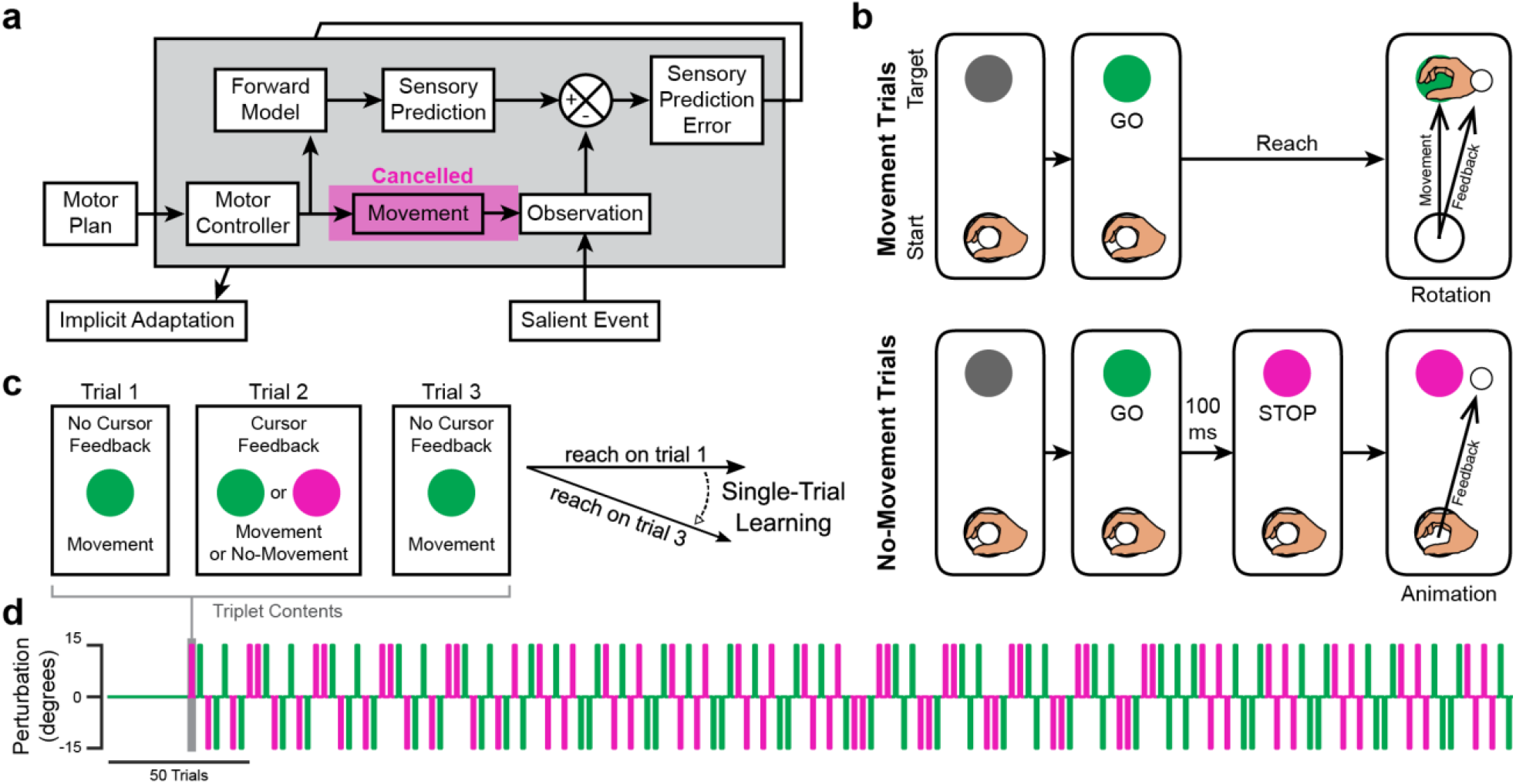
Schematics showing the proposed learning framework and task design. (**a**) Schematic showing how the forward model may support implicit motor adaptation in the presence of sensory feedback not causally related to self-generated movement. (**b**) Events on trials with visual feedback. The robotic apparatus brought the participant’s hand to the starting location to initiate a trial. On Movement trials (top), the target turned green (GO), cueing participants to reach through the target. On trials with visual feedback, participants observed a white feedback cursor move along a rotated trajectory (Rotation). On No-Movement trials (bottom), the target turned magenta 100 ms after turning green, cueing participants to withhold movement (STOP). After a delay, an animation showing the feedback cursor moving 15° off-target played (Animation). The hand is shown in the figure for illustrative purposes but was not visible during the experiment. (**c**) How single-trial learning (STL) was computed using a triplet paradigm. Triplets were composed of 2 Movement trials without visual feedback flanking either a Movement or a No-Movement trial with visual feedback. STL was measured as the difference between reach angles on the flanking trials. (**d**) The pseudorandomized order in which trials were presented for an example participant. Color indicates movement condition (Movement: green, No-Movement: magenta).

Prior work has illustrated that higher-level cognitive processes support visuomotor learning without movement, for instance when observers witness others’ motor errors: motor learning in this case might be driven by SPEs, or by other types of performance errors beyond SPE (e.g., reward prediction errors), or perhaps by a combination of multiple sources of error (14–16). Here, we isolated implicit motor adaptation to specifically test whether SPE computation requires movement execution, as SPEs are both necessary and sufficient for this form of learning (17–23). Having isolated implicit motor adaptation, we then asked whether withheld movements could undergo adaptation following the observation of “simulated” sensory feedback.

To that end, we measured trial-by-trial implicit adaptation during a visuomotor task in which human participants saw visual feedback while performing – or withholding – hand and arm movements, using a modified stop-signal paradigm. To isolate implicit adaptation, we employed a recently-developed approach that requires participants to direct their movements directly for presented targets and disregard visual feedback (22, 24–28). We predicted that single-trial motor adaptation would occur following both typical movement trials that generated sensory error, as well as trials where movements were withheld but simulated sensory errors were observed. If confirmed, this result would demonstrate that the brain can compute SPEs in the absence of movement and can thus drive the adaptation of planned movements that were never performed.

## RESULTS

### Simulated and typical visuomotor rotations cause motor adaptation

In our first experiment, we measured implicit motor adaptation in humans (*n* = 20) performing or withholding straight reaches during a visuomotor adaptation task (**Fig. 1b**). Vision of the hand and arm was occluded by a mirror that reflected visual feedback from a horizontally mounted monitor. A white cursor provided feedback about participants’ hand positions as they reached from a starting location to a displayed target. After a brief acclimation period, trials were organized into triplets, such that each trial with cursor feedback was flanked by trials without cursor feedback. This allowed for a reliable measurement of single-trial learning (STL) in response to feedback, quantified as the difference between the direction of hand movement (hand angle) on the first and third trials of each triplet (**Fig. 1c**). Trials with cursor feedback were either Movement trials on which a Go signal prompted movement or No-Movement trials on which a Stop signal immediately followed the Go signal, indicating that movements should be withheld. On Movement trials, feedback involved a visuomotor error (±15° rotation added to the visual cursor path; + = clockwise; **Fig. 1b right**). On No-Movement trials, sensory feedback involved a simulation of the cursor’s path, using timing variables based on ongoing measurements of participant behavior (see *Methods*). All flanking trials of each triplet were Go trials and required movements. The direction of the error (clockwise [CW] or counterclockwise [CCW]) was pseudorandomly varied across triplets to maintain overall adaptation near 0 throughout the session (**Fig. 1d**). This straightforward design allowed us to test the hypothesis that SPE computation and motor adaptation do not require that movement and sensory feedback to be causally linked (**Fig. 1a**).

Consistent with our predictions, rotated cursor paths on Movement and No-Movement trials both caused subsequent hand trajectories to shift opposite the direction of the rotation (**Fig. 2b-c**), with a 2-way repeated measures ANOVA revealing statistically significant main effects of the direction (CW vs CCW) of the perturbation applied (*F*(1, 19) = 98.62, *p* = 5.89 × 10^−9^, 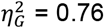). While there was no significant main effect of withholding movement (*F*(1, 19) = 1.79, *p* = 0.20), we observed a significant interaction between the perturbation applied and withholding movement (*F*(1, 19) = 137.32, *p* = 3.87 × 10^−10^, 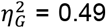). Post-hoc pairwise comparisons confirmed that STL was sensitive to perturbation direction during both Movement (paired t-test: *t*(19) = 12.92, *p*_*adj*_ = 2.96 × 10^−10^, *Cohen’s d* = 5.12) and No-Movement triplets (*t*(19) = 4.39, *p*_*adj*_ = 3.13 × 10^−4^, *Cohen’s d* = 1.63), and also indicated that STL magnitude was greater across Movement than No-Movement triplets (paired-samples signed-rank test, CW rotations: *V* = 210, *p*_*adj*_ = 2.55 × 10^−6^, *r* = 0.88; CCW rotations: *t*(19) = 9.43, *p*_*adj*_ = 2.70 × 10^−8^, *Cohen’s d* = 2.02) rotations. The overall amplitude of adaptation observed both with and without movement was within the range of implicit learning rates measured in previous studies (**Supplemental Fig. 1**).

**Figure 2.**
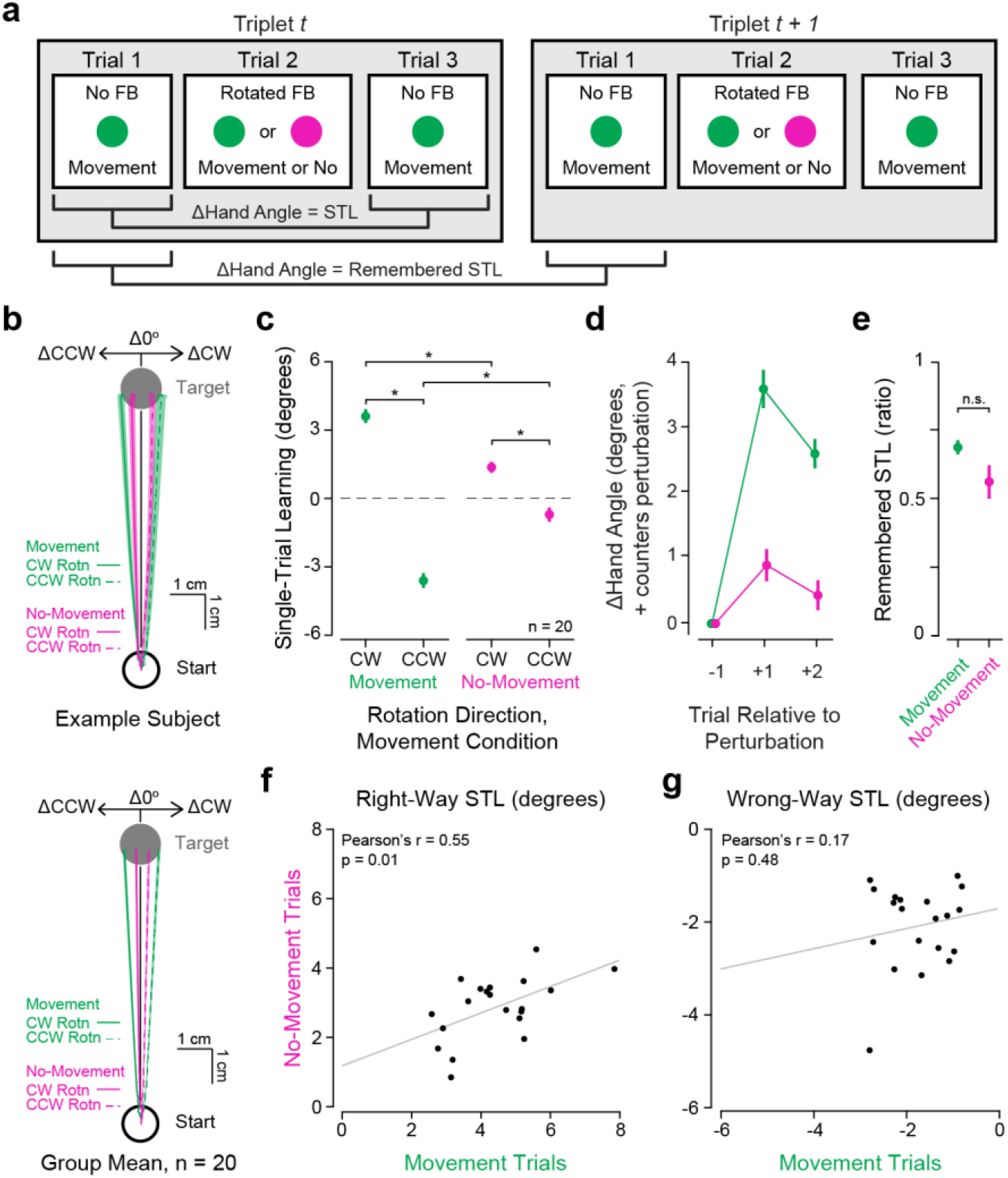
Effects of typical and simulated errors during a visuomotor reach adaptation task. (**A**) Schematic illustrating how STL and remembered STL measurements were computed. (**B**) An example subject’s (top) and the group’s (bottom) mean ± SEM changes in reach paths across triplets with a rotation applied (green: triplets with perturbations on Movement trials, magenta: triplets with perturbations on No-Movement trials, solid lines: perturbation was a CW rotation, dashed lines: perturbation was a CCW rotation). (**C**) STL across Movement (green) and No-Movement (magenta) triplets for all participants (*n* = 20). Positive changes in hand angle are CCW. Refer to Supplemental Table 1 for details on all statistical tests. (**D**) Group mean ± SEM Δhand angle values after exposure to Movement (green) and No-Movement (magenta) trial perturbations. Positive Δ values indicate that the change in hand angle proceeded opposite the direction of the perturbation (*i.e*., the direction that would counter the error, “Right-Way”). (**E**) Group mean of participants’ ratios of remembered STL to initial STL during Movement and No-Movement trials. (**F**) The relationship between Right-Way STL observed during Movement and No-Movement triplets. (**G**) As in (**F**), but for STL observed on trials where adaptation proceeded in the direction that would exacerbate the error (*i.e*., the same direction as the perturbation applied, “Wrong Way”). Statistical significance (* = *p*_*adj*_ < 0.05; n.s. = *p*_*adj*_ ≥ 0.05) is indicated. Abbreviations: STL – single-trial learning, CW – clockwise, CCW – counterclockwise, Δ – change in.

**Supplemental Figure 1.**
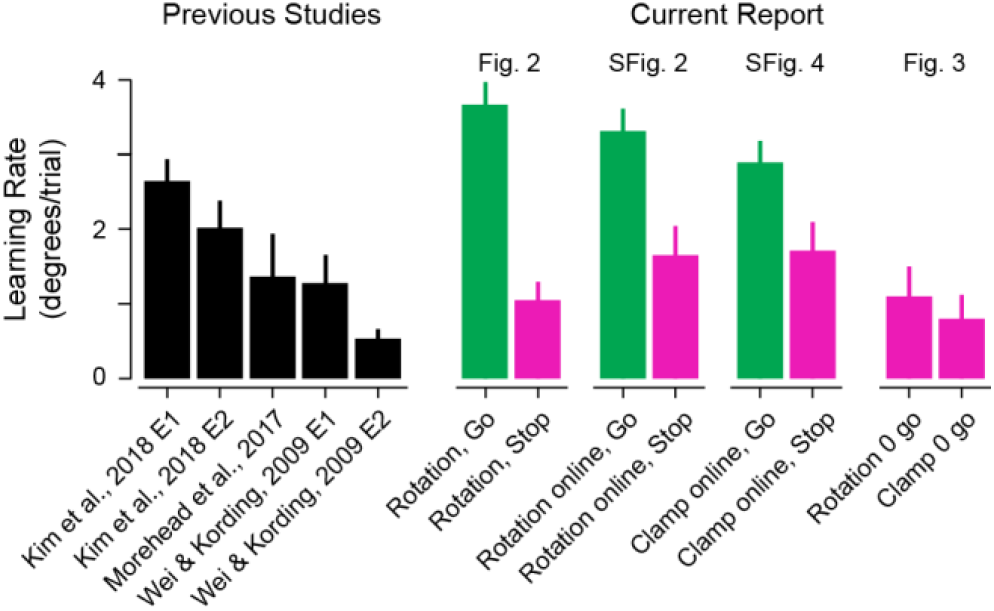
Learning rates reported in the literature and observed in the current study. Learning rates for motor adaptation observed in previous studies are shown at left in black, and learning rates observed in each experiment in the current report are shown at the right, with data from Movement triplets shown in green and data from No-Movement triplets shown in magenta. Data are shown as mean ± SEM, and are shown for rotational/error clamp perturbations of 15°, with the exception of Wei & Kording, 2009 E2, where an 11° perturbation was applied. Papers referred to and their corresponding reference numbers: Kim et al., 2018 (25); Morehead et al., 2017 (22); Wei & Körding, 2009 (29). “Rotation, Go” and “Rotation, Stop” show data from the in-lab experiment where participants saw 15° rotated feedback on Movement trials (i.e., data from Fig. 2), “Rotation online, Go” and “Rotation online, Stop” show data from the online experiment where participants saw 0-15° rotated feedback on Movement trials (i.e., data from Supplemental Fig. 2). “Clamp online, Go” and “Clamp online, Stop” show data from the online experiment where participants saw 0-15° error-clamed feedback (i.e., data from Supplemental Fig., 4). “Rotation 0 go” and “Clamp 0 go” show data from the online experiments where participants saw 0° perturbed feedback on Movement trials. Abbreviations: E, experiment.

To address whether observed STL measured genuine implicit learning, we checked whether adaptation persisted beyond the trial after an error was experienced. We examined participants’ hand angles on the second trial after a perturbation relative to the pre-perturbation baseline trial (i.e., hand angle on trial 1 of triplet *t + 1* relative to hand angle on trial 1 of triplet *t*, subsequently referred to as remembered STL, **Fig. 2a**). As visual feedback was withheld on both trial types, this approach provided a pure measure of persistent memory in the absence of error-driven changes in performance. Hand angle remained adapted in the direction opposite the rotation on trials with nonzero perturbations regardless of movement condition (**Fig. 2d**), suggesting that genuine implicit learning was observed in response to errors under both movement conditions. Closer examination of the relative ratio of remembered STL to initial STL revealed that retention of adaptation differed significantly from zero after both Movement (*t*(19) = 26.20, *p*_*adj*_ = 6.71 × 10^−16^, *Cohen’s d* = 5.86) and No-Movement triplets (*t*(19) = 9.20, *p*_*adj*_ = 2.98 × 10^−8^, *Cohen’s d* = 2.06), and the amount of retention observed was not statistically significantly different between the movement conditions (*t*(19) = 2.07, *p*_*adj*_ = 0.053, **Fig. 2e**).

To assess the potential similarity of mechanisms underlying adaptation after errors on Movement and No-Movement trials, we compared STL amplitude under each condition, reasoning that there should be a reliable relationship between the two measures if STL is supported by the same mechanism following both Movement and No-Movement trials. When we considered instances of STL in the direction that would compensate for the observed error (the direction opposite the rotation, i.e., the “Right-Way”), within-subject changes in hand angle were correlated between Movement and No-Movement trials (*Pearson’s r* = 0.55, *p* = 0.01; **Fig. 2f**). Conversely, changes in hand angle in the direction that would exacerbate the observed error (the direction of the rotation, i.e., the “Wrong-Way”) were uncorrelated between Movement and No-Movement trials (*Pearson’s r* = 0.17, *p* = 0.48, **Fig. 2g**). Together, these observations suggest that the same learning process may underlie adaptive STL events in response to errors during both kinds of trials, while maladaptive changes in hand angle may be attributable to noise.

### Implicit motor adaptation proceeds after simulated errors in an online visuomotor task

Illustrating that the above observations are reproducible and generalize across experimental contexts, we again observed that simulated errors in No-Movement trials also induced motor adaptation in an online, crowd-sourced version of the task. Participants (*n* = 40) made hand movements using their computer mouse or trackpad to move a cursor towards a target. As in the experiment described above, trials were presented in triplets, allowing us to measure STL in response to cursor feedback presented during Movement and No-Movement trials at the center of each triplet (**Fig. 1b-d**). For this online study, triplets with 0° perturbations/simulated errors were also included to provide an estimate of baseline changes in hand angle, in the event that participants exhibited strong movement biases in the online platform.

STL was directionally appropriate for the perturbation applied during both Movement and No-Movement trials (**Supplemental Fig. 2a-c**, please refer to the supplemental material details of the statistical analysis). Further echoing the results of the in-person study, STL on both Movement and No-Movement trials was retained beyond the triplet in which the relevant error occurred (**Supplemental Fig. 2d-e**), and the magnitude of STL in the direction that would counter the perturbation was again correlated across the two movement conditions (**Supplemental Fig. 2e**). These data provide further support for the claim that movements that are not performed can undergo implicit motor adaptation, and they extend our findings to a task with different movement demands (e.g., finger or wrist movements versus full, center-out reaches).

**Supplemental Figure 2.**
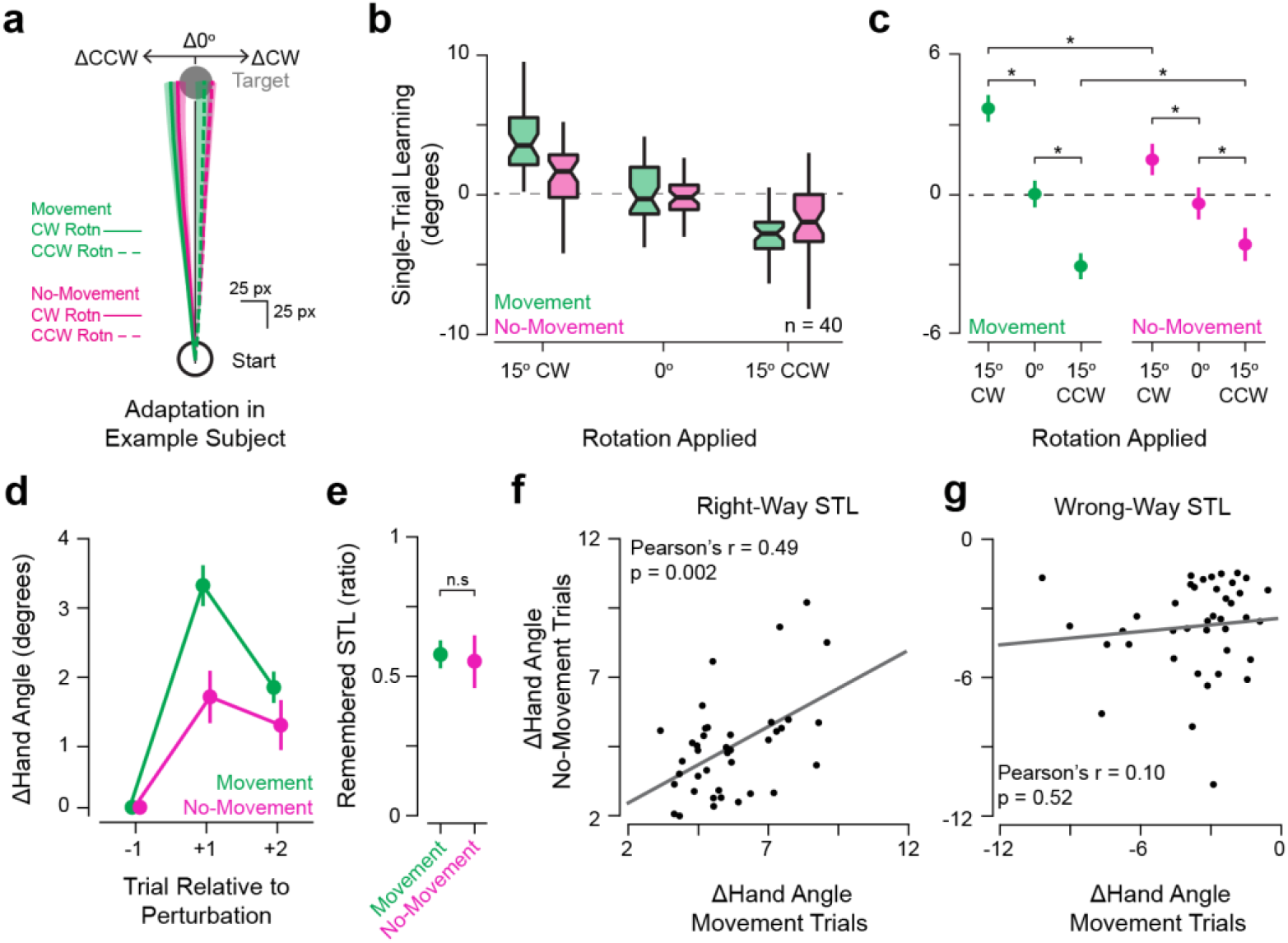
Single-trial learning in response to errors on Movement and No-Movement trials during an online visuomotor adaptation task. (**a**) An example participant’s mean ± SEM changes in reach paths across triplets (green: triplets with perturbations on Movement trials, magenta: triplets with perturbations on No-Movement trials, solid lines: perturbation was a CW rotation, dashed lines: perturbation was a CCW rotation). (**b**) Boxplot showing STL across Movement (green) and No-Movement (magenta) triplets for participants in an online version of the task described in Figure 1 (*n* = 40). (**c**) Estimated marginal means (EMMs) ± 95% confidence intervals from the linear mixed model (LMM) fit to participants’ STL performance (summarized in **b**). The LMM (fixed effects: rotation [15° counterclockwise {CCW}, 0°, and 15° clockwise {CW}], movement condition [Movement, No-Movement], rotation x movement condition interaction; random effects: participant) revealed significant main effects of rotated cursor feedback (*F*(2, 2223) = 136.46, *p* = 2.2 × 10^−16^, *partial R*^*2*^ = 0.11) and movement condition (*F*(1, 2248) = 4.74, *p* = 0.03, *partial R*^*2*^ = 0.002), as well as a significant interaction (*F*(2, 2229) = 12.40, *p* = 4.41 × 10^−6^, *partial R*^*2*^ = 0.01). Post-hoc pairwise comparisons of the EMMs from the model support the claim that rotated feedback induced a statistically significant degree of STL on both Movement (0° vs 15° CW: *t*(2227) = 9.14, *p*_*adj*_ = 6.39 × 10^−19^, *Cohen’s d* = 0.61; 0° vs 15° CCW: *t*(2220) = 7.81, *p*_*adj*_ = 2.61 × 10^−14^, *Cohen’s d* = 0.52) and No-Movement trials (0° vs 15° CW: *t*(2225) = 3.92, *p*_*adj*_ = 1.39 × 10^−4^, *Cohen’s d* = 0.31; 0° vs 15° CCW: *t*(2229) = 3.56, *p*_*adj*_ = 4.84 × 10^−4^, *Cohen’s d* = 0.29). Adaptation in the presence of a rotation was significantly greater in Movement trials than No-Movement trials for CW (*t*(2238) = 4.98, *p*_*adj*_ = 1.26 × 10^−6^, *Cohen’s d* = 0.37) and CCW rotations (*t*(2239) = 2.06, *p*_*adj*_ = 0.04, *Cohen’s d* = 0.15). (**d**) Group mean ± SEM change in (Δ) hand angle after exposure to Movement (green) and No-Movement (magenta) triplets’ perturbations. Positive Δ values indicate that the change in hand angle proceeded opposite the direction of the perturbation (*i.e*., in the direction that would counter the error). (**e**) Group mean ± SEM ratio of remembered STL to STL. Remembered STL was statistically significantly greater than 0 for both Movement (one-sample signed-rank test: *V* = 819, *p*_*adj*_ = 1.09 × 10^−11^, *r* = 0.87) and No-Movement triplets (*V* = 769, *p*_*adj*_ = 9.69 × 10^−s8^, *r* = 0.76), but remembered STL did not significantly differ between movement conditions (paired-samples signed-rank test: *V* = 441, *p*_*adj*_ = 0.68). (**f**) Scatter plot showing the relationship between individual subjects’ STL amplitude in the direction opposite the rotation on Movement and No-Movement trials. When we considered instances of STL in the direction that would compensate for the observed error (update opposite rotation, “Right-Way”), within-subject changes in hand angle were correlated between Movement and No-Movement trials (*Pearson’s r* = 0.49, *p*_*adj*_ = 0.002). (**g**) As in **f**, but showing data from trials with changes in hand angle in the direction that would exacerbate the observed error (update in direction of rotation, “Wrong-Way”). These ΔHand Angle values were uncorrelated between Movement and No-Movement trials (*Pearson’s r* = 0.10, *p*_*adj*_ = 0.52). These observations support the idea that the same learning process may underlie adaptive single-trial learning events in response to errors on both kinds of trials, while maladaptive changes in hand angle may be attributable to potential sources of random noise. Boxplot centers: median, notch: 95% confidence interval of the median, box edges: 1^st^ and 3^rd^ quartiles, whiskers: most extreme value within 1.5*interquartile range of the median. Statistical significance (* = p < 0.05; n.s. = p ≥ 0.05) is. Abbreviations: STL – single-trial learning, Δ – change, CW – clockwise, CCW – counterclockwise.

### Motor adaptation during No-Movement triplets does not depend on participants’ control over cursor trajectory during Movement trials

We note that rotated visual feedback on Movement trials was sensitive to people’s actual reaching directions because the rotation was simply added to the measured reach direction, as is typical in visuomotor rotation tasks. It is possible that these directional contingencies affected participants’ responses to error, potentially encouraging them to attempt to deliberately control the cursor’s position via an explicit re-aiming process (23). To rule this out, we recruited a new group of participants (*n* = 37) to perform a variant of the task where the visual cursor moved in a fixed path (“error-clamped” feedback (22); **Supplemental Fig. 3a**) in one of three directions (0° or 15° CW/CCW) on the trials with feedback.

**Figure 3.**
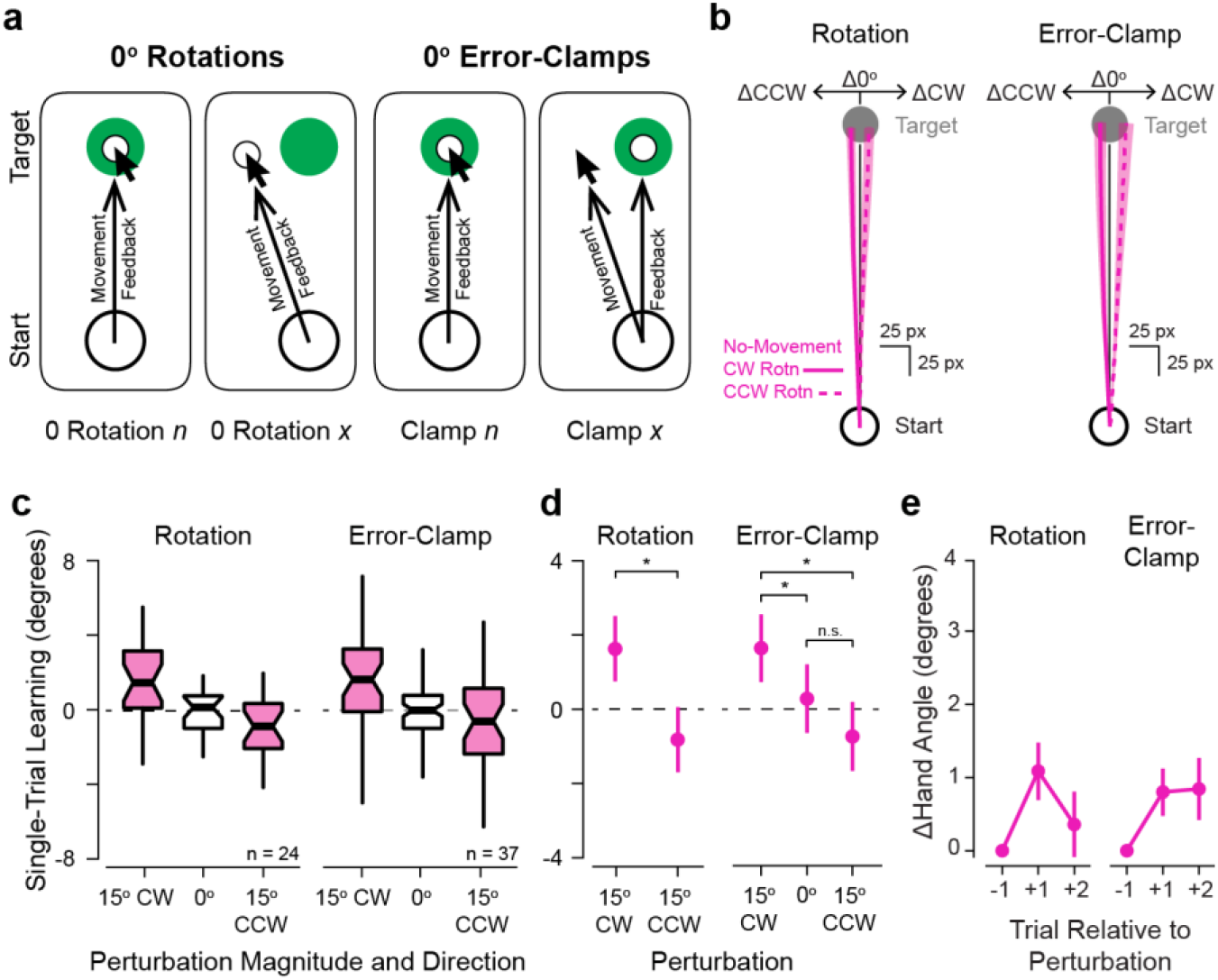
Effects of simulated errors when perturbations were never applied during Movement trials. (**a**) Schematic illustrating the relationship between movement and visual feedback on Movement trials during an experiment where visuomotor rotations (left) or error-clamps (right) were never applied during Movement trials. (**b**) An example participant’s mean ± SEM changes in reach paths across No-Movement triplets from studies in which non-zero rotations (left) and error-clamps (right) were never applied (solid lines: perturbation was CW, dashed lines: perturbation was CCW). (**c**) Boxplots showing STL in response to different directions of simulated errors (No-Movement triplets indicated in magenta) from rotation (left, *n* = 24) and error-clamp (right, *n* = 37) studies. (**d**) Estimated marginal means ± 95% confidence intervals from the linear mixed models fit to each participant’s STL performance summarized in (**c**). Asterisks indicate statistically significant differences. (**e**) Mean ± SEM relative hand angles on the two trials after a perturbation was presented on a No-Movement trial. Please refer to Supplemental Table 3 for detailed statistical results. Boxplot centers: median, notches: 95% confidence interval of the median, box edges: 1^st^ and 3^rd^ quartiles, whiskers: most extreme values within 1.5*IQR of the median. Statistical significance (* = *p*_*adj*_ < 0.05; n.s. = *p*_*adj*_ ≥ 0.05) is indicated for selected comparisons. Abbreviations: STL – single-trial learning, CW – clockwise, CCW – counterclockwise, Δ – change in.

Replicating and extending the findings reported above, participants assigned to the error-clamp condition exhibited STL after Movement and No-Movement trials (**Supplemental Fig. 3b-d**; please refer to supplemental material for details). We also observed significant retention of STL (**Supplemental Fig. 3e-f**) and a significant correlation between STL amplitude on Movement and No-Movement trials where adaptation proceeded opposite the direction of the perturbation (**Supplemental Fig. 3g-h**). These data further strengthen the claim that motor adaptation does not require movement.

**Supplemental Figure 3.**
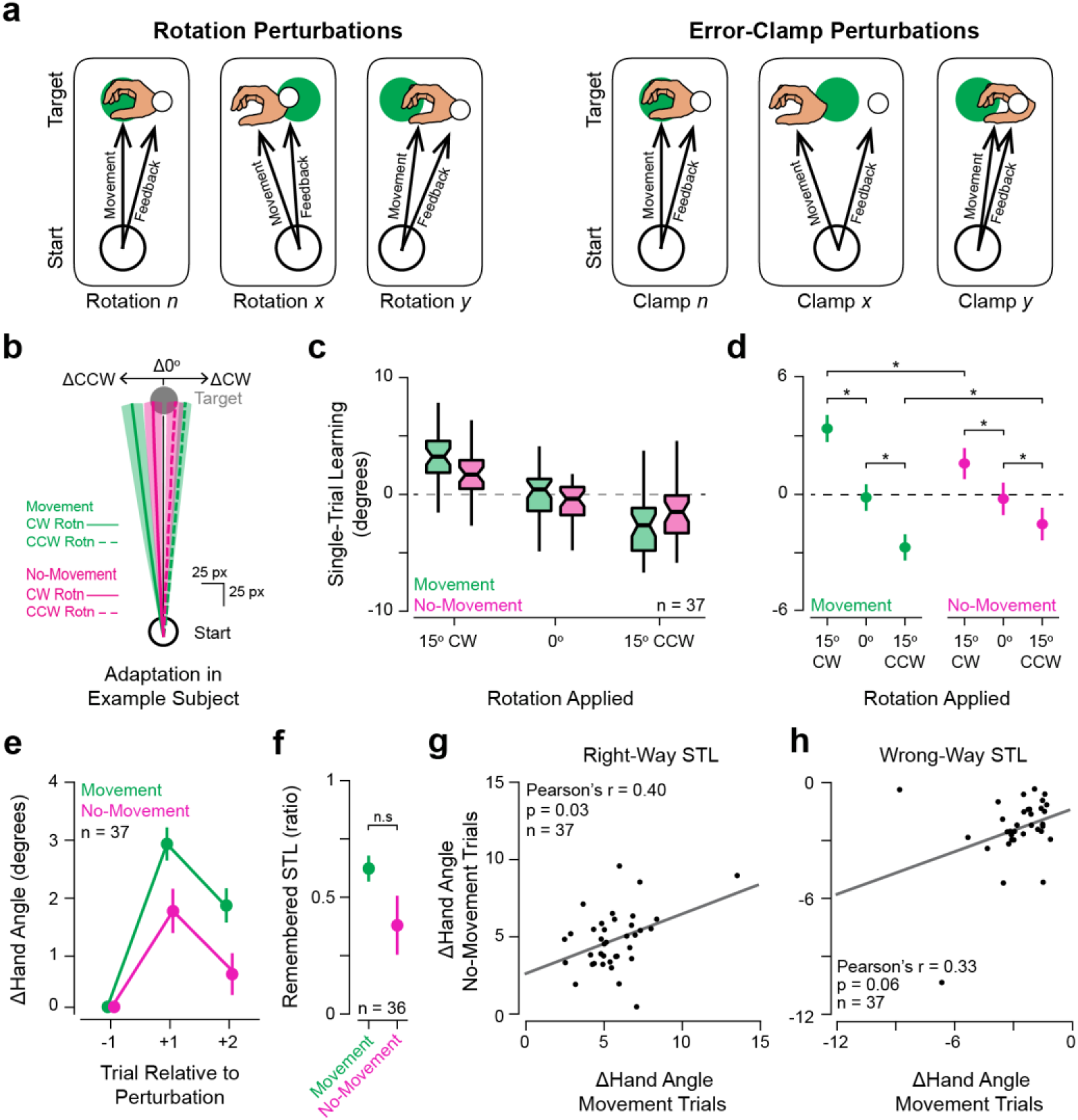
Single-trial learning in response to errors on Movement and No-Movement trials with error-clamped feedback or simulated errors. (**a**) Diagrams showing the relationship between hand and cursor feedback movement directions under rotational (left) and error-clamp regimes (right). When rotations are applied, the cursor’s movement direction is contingent upon the participant’s movement direction. When error-clamp perturbations are applied, the cursor travels in a fixed direction, regardless of the direction that the hand travels. As error-clamp perturbations render deliberate changes in movement direction useless, they are often used in studies attempting to isolate implicit motor adaptation processes. (**b**) An example participant’s mean ± SEM changes in reach paths across triplets (green: triplets with perturbations on Movement trials, magenta: triplets with perturbations on No-Movement trials, solid lines: perturbation was CW error-clamp, dashed lines: perturbation was CCW error-clamp). (**c**) Boxplot showing STL across Movement (green) and No-Movement (magenta) triplets for participants (*n* = 37) in an online experiment where cursor feedback was error-clamped on Movement trials. (**d**) Estimated marginal means (EMMs) ± 95% confidence intervals from the linear mixed model (LMM) fit to participants’ STL performance (summarized in **c**). The LMM (fixed effects: rotation [15° counterclockwise {CCW}, 0°, and 15° clockwise {CW}], movement condition [Movement, No-Movement], error-clamp x movement condition interaction; random effects: participant) revealed significant main effects of error-clamped cursor feedback (*F*(2, 1829) = 79.46, *p* = 2.2 × 10^−16^, *partial R*^*2*^ = 0.08) and an interaction between error-clamp and movement condition (*F*(2, 1832) = 8.45, *p* = 0.0002, *partial R*^*2*^ = 0.0003), although there was no main effect of movement condition (*F*(1, 1844) = 0.60, *p* = 0.44). Post-hoc comparisons of the EMMs from the model revealed significant STL in response to non-zero error-clamped feedback on both Movement (0° vs 15° CW: *t*(1827) = 7.55, *p*_*adj*_ = 3.08 × 10^−13^, *Cohen’s d* = 0.56; 0° vs 15° CCW: *t*(1828) = 5.57, *p*_*adj*_ = 8.84 × 10^−8^, *Cohen’s d* = 0.41) and No-Movement trials (0° vs 15° CW: *t*(1830) = 3.21, *p*_*adj*_ = 0.002, *Cohen’s d* = 0.29; 0° vs 15° CCW: *t*(1832) = 2.25, *p*_*adj*_ = 0.03, *Cohen’s d* = 0.22). Adaptation in the presence of a 15° error-clamp was significantly greater on Movement trials than No-Movement trials for CW (*t*(1846) = 3.49, *p*_*adj*_ = 0.0009, *Cohen’s d* = 0.29) and CCW clamps (*t*(1846) = 2.29, *p*_*adj*_ = 0.03, *Cohen’s d* = 0.19). Please refer to Supplementary Table 2 for further details on post-hoc comparisons in this panel. (**e**) Group mean ± SEM change in (Δ) hand angle one and two trials after exposure to Movement (green) and No-Movement (magenta) triplets’ perturbations. Positive Δ values indicate that the change in hand angle proceeded opposite the direction of the perturbation (*i.e*., in the direction that would counter the error). (**f**) Remembered STL shown as the ratio of relative hand angle 2 trials after experiencing a perturbation to the relative hand angle 1 trial after the perturbation (STL). Remembered STL was significantly greater than 0 after both Movement (green; one-sample t-test: *t*(36) = 11.31, *p*_*adj*_ = 6.23 × 10^−13^, *Cohen’s d* = 1.86) and No-Movement triplets (magenta, one-sample signed-rank test: *V* = 579, *p*_*adj*_ = 5.95 × 10^−5^, *r* = 0.64), but did not exhibit statistically significant differences between movement conditions (paired t-test: *t*(35) = 1.71, *p*_*adj*_ = 0.09). Remembered STL on No-Movement trials could not be computed for one participant, so *n* = 36 instead of 37 in this panel. (**g**) Scatter plot showing the relationship between individual subjects’ STL amplitude in the direction opposite the error-clamp on Movement and No-Movement trials (i.e., the “Right-Way”). Right-way changes in hand angle were correlated between Movement and No-Movement trials (*Pearson’s r* = 0.40, *p*_*adj*_ = 0.03). (**h**) As in (**g**), but showing data from trials on which STL proceeded in the same direction as the error-clamp (*i.e*., the “Wrong-Way”). Wrong-Way changes in hand angle were not statistically significantly correlated between Movement and No-Movement trials (*r* = 0.33, *p*_*adj*_ = 0.06). Boxplot centers: median, notch: 95% confidence interval of the median, box edges: 1^st^ and 3^rd^ quartiles, whiskers: most extreme value within 1.5*interquartile range of the median. Statistical significance (* = p < 0.05; n.s. = p ≥ 0.05) is indicated for selected comparisons. Abbreviations: STL – single-trial learning, Δ – change, CW – clockwise, CCW – counterclockwise.

### Adaptation during No-Movement triplets does not depend on within-session adaptation during Movement triplets

In two further experiments, we asked if adaptation to errors in the No-Movement condition was contingent on sharing a context with the Movement condition. In other words, if learning in the No-Movement condition only occurs when there are neighboring trials in the Movement condition producing typical SPEs, it is possible that adaptive responses observed in the No-Movement condition reflect a “cueing” effect, whereby an adapted sensorimotor map is cued by observation of the visual error and then retrieved on the subsequent trial(s) (30, 31). While our previous retention (**Fig. 2d-e, Supplemental Fig. 2d-e, Supplemental Fig. 3e-f**) and correlation (**Fig. 2f-g, Supplemental Fig. 2f-g, Supplemental Fig. 3g-h**) results argue against this interpretation as they suggest a shared learning mechanism across movement conditions, we opted to directly test this alternative explanation in another pair of experiments. Here, we only included 0° rotated (**Fig. 3a**, left, *n* = 24 participants) or clamped (**Fig. 3a**, right, *n* = 37 participants) error feedback on Movement trials, but maintained 0° or 15° CW/CCW errors on the No-Movement trials. Thus, visual perturbations were never paired with movement. The key results were again replicated – learning was preserved in the No-Movement condition even when error feedback had never been associated with executed movements (**Fig. 3c**, rotation: LMM: *F*(557) 23.01, *p* = 2.07 × 10^−6^, *partial R*^*2*^ = 0.04; error-clamp: *F*(802) = 9.41, *p* = 9.14 × 10^−5^, *partial R*^*2*^ = 0.02). Post-hoc pairwise comparisons showed that adaptation was significantly different between triplets with clockwise and counterclockwise errors for both the rotation (*t*(557) = 4.80, *p* = 2.07 × 10^−6^, *Cohen’s d* = 0.4) and error-clamp experiments (*t*(1453) = 4.32, *p* = 5.34 × 10^−5^, *Cohen’s d* = 0.37) – a hallmark of implicit motor adaptation (please refer to **Supplemental Table 3** for all post-hoc test results). Overall levels of STL observed on No-Movement trials were comparable during these two experiments to those discussed above, and within the range of learning rates previously observed in the literature (**Supplemental Fig. 1**). Furthermore, both groups of participants showed retention of STL that differed significantly from 0 (rotation, mean ± SEM: 0.53 ± 0.06 retention ratio, one-sample t-test: *t*(22) = 8.28, *p* = 3.34 × 10^−8^, *Cohen’s d* = 1.73; error-clamp, median: 0.45, interquartile-range: 0.58, one-sample signed-rank test: *V* = 507, *p* = 0.001, *r* = 0.53). Overall, these experiments support the hypothesis that motor adaptation can proceed without movement execution.

## DISCUSSION

Our results demonstrate that movements can be implicitly refined even when they are not performed. Participants who were cued to reach towards a target but suppressed that movement after observation of a Stop cue showed consistent, robust STL in response to simulated errors (**Figs. 2-3, Supplemental Fig. 2, Supplemental Fig. 3**). As implicit learning necessarily proceeds following SPEs, our data also provide evidence that SPEs are computed even when movements are not performed. These findings strongly support the fundamental assumptions of predictive processing frameworks of motor adaptation, where precise sensory predictions are generated from a movement intent (or “plan”, “goal”) and compared against sensory observations to induce error-based learning (8, 11, 13, 32, 33).

We argue that we have measured learning via an implicit process, and, by extension, that the STL observed in our study provides evidence that SPEs are computed regardless of whether a movement is performed. Although visuomotor tasks sometimes recruit cognitive strategies (e.g., deliberate “re-aiming” of movements), multiple factors indicate that our studies successfully measured implicit adaptation (23, 34, 35). First, participants were instructed to ignore the displayed cursor and try to contact the target on every trial, a straightforward technique which has been consistently shown to eliminate the explicit re-aiming of movements (20, 22, 24–26, 28). Second, randomization of the presence and direction of errors discourages explicit learning, reducing motivation to apply ineffective re-aiming strategies (see (36)). Third, data from participants who did appeared to not fully recall the instruction to always aim directly at the target were excluded (see *Methods*), decreasing the likelihood that strategic re-aiming contaminated the analysis. Fourth, adaptation persisted into subsequent no-feedback trials (**Fig. 2d-e, Fig. 3e, Supplemental Fig. 2d-e, Supplemental Fig. 3e-f**), consistent with lingering implicit motor learning; it is unlikely that strategies would be maintained through trials where no feedback is expected. Fifth, the magnitude of STL observed was generally consistent with multiple previous studies that similarly measured implicit motor adaptation rates (**Supplemental Fig. 1**) (22, 25, 37). Lastly, the adaptation effects observed in the No-Movement conditions were not attributable to the recall of learning that had occurred on Movement trials (**Fig. 3**). Our data thus provide converging evidence that movement is not required for implicit adaptation, and, by extension, SPE computation.

While motor planning and concurrent sensory observations are sufficient to drive SPE computation and motor adaptation, our data also indicate that participants showed significantly stronger STL over triplets with Movement trials versus No-Movement trials. This suggests that movement provides additional training input to the brain. Interestingly, this is consistent with patterns of cerebellar activity during motor behaviors, and current thinking about mechanisms for learning in cerebellar-dependent tasks like implicit reach adaptation (22). Purkinje cell complex spikes are a powerful teaching signal in the cerebellum, and these complex spikes exhibit firing patterns that may be movement-dependent (38–41). During target-directed reaching, complex spikes related to reach goal locations are generated *after* reach onset (42). If these complex spikes are tied to motor performance and not motor planning, then the absence of these error signals on No-Movement trials may account for reduced levels of STL without movement (43–46). Another non-mutually exclusive possibility is that the precise timing of SPEs is less effective in our No-Movement condition than under normal movement conditions: in the former case, the timing of simulated feedback is controlled by the experimenter and not triggered by the subject’s actual movement, potentially adding a novel source of noise into the adaptation process (47, 48). Irrespective of the fact that STL was of lesser amplitude across No-Movement than Movement triplets, our data demonstrate the significant influence of the brain’s prediction signals on learning – even without the ability to directly attribute sensory feedback to an actual movement, prediction of a planned movement’s sensory consequences supports the error computations that drive adaptation of future behavior.

Our findings add to a body of work indicating that many forms of motor learning do not strictly require movement-based practice. For instance, in (49), after human participants observed others adapting to a force field applied during reaching movements, the observers were able to partially compensate for that same force field when they encountered it themselves. Interestingly, this observational learning did not proceed if participants were executing other task-irrelevant movements during the observation period. This finding has been linked to subsequent neuroimaging data showing that observational learning recruits brain areas associated with motor planning, and together are taken to suggest that engagement in a covert motor planning process may allow for force-field adaptation via observation (49–51). Together, this related prior work and the evidence we have provided here suggest that there may be multiple routes to inducing motor planning and ultimately driving motor adaptation.

Other reports in the motor learning literature have provided evidence for cognitive compensation for observed motor errors during reaching, improved visual tracking following observation of target movement without engagement in visual pursuit, and improvement in movement speeds as a result of mental imagery training; this work highlights the breadth of motor performance-related processes that can be trained without engagement in physical movements (14, 15, 52, 53). Together with the findings related to motor adaptation via observation discussed above, the findings of the present report suggest that many features of motor performance can be improved by training regiments that do not involve movement. This points to a potential opportunity for the development of motor training or rehabilitation protocols that can be used when people are unable to physically perform target motor behaviors, perhaps improving performance beyond what physical practice can do alone.

Finally, our results echo the fact that other types of learning can occur without overt task execution. As an example, fear associations can be extinguished by instructing participants to imagine a fear-predicting stimulus even when they are not presented with the stimulus, with this “imagination” protocol generating neural signatures of the negative prediction errors observed during naturalistic fear extinction (54, 55). Considering both this prior work and the findings presented in this study, it may be that the generation of predictions for comparison with sensory observations is sufficient for error-based learning across motor and non-motor domains alike. In other words, task execution may not always be required for learning, so long as the predictions and observations needed to compute errors are both present.

## ACKNOWLEDGEMENTS

This work was supported by a grant from the National Institutes of Health to OK (F32-NS122921).

## METHODS

### Participants

Participants (*n* = 233, aged 18-35, 126 female) recruited from the research participation pools at Princeton/Yale University and on Prolific provided informed consent, approved by each University’s IRB. Seventy-five participants were excluded (10 from the dataset collected for Supplemental Fig. 2, 13 from the dataset collected for Supplemental Fig. 3, 26 from the dataset collected for the rotation perturbation experiments described in Fig. 3, and 26 from the dataset collected for the error-clamp perturbation experiments described in Fig. 3) for failure to sufficiently recall task instructions, as ascertained by a questionnaire at the end of the experiment, leaving 158 participants for our analyses. See the *Questionnaire* section below the *Test phase* sections for more details. We note that all the key results described here (*i.e*., statistically significant learning after No-Movement trials) held with or without these exclusions; we opted to be conservative in our exclusion criteria to limit potential effects of explicit learning. We note that the samples used for the online experiments described in the text are around twice the size of similar studies in the literature, providing additional statistical power to compensate for the experiment being conducted remotely.(56–58)

### Task Setup: In Lab

Participants were seated in a chair and made ballistic reaching movements while grasping the handle of a robotic manipulandum with their dominant hand (Kinarm End-Point). The manipulandum restricted movements to the horizontal plane. All visual stimuli were projected to the participant via a horizontal display screen (60 Hz) reflected onto a semi-silvered mirror mounted above the robotic handle. The mirror occluded vision of the arm, hand, and robotic handle, preventing direct visual feedback of hand position. Tasks were programmed in Matlab 2019a’s Simulink for deployment in Kinarm’s Dexterit-E software (version 3.9). Movement kinematics were recorded at 1 kHz. Each participant viewed a single target located at either 45°, 135°, 225°, or 315° (with target position counterbalanced across participants), 8 cm from a central starting location. The target was visible throughout the experiment.

### Task Setup: Online

Experiments were conducted remotely using a custom JavaScript web application based on Phaser 3.24 (download available at (59)), similar to an approach previously described.(60) Each participant viewed a single target located at either 45°, 135°, 225°, or 315° (with target position counterbalanced across participants), 250 pixels from a central starting location. The target was visible throughout the experiment.

Participants used an input device of their choice to control their computer cursor during center-out movements. One participant reported using a touchscreen device and was excluded from all analyses. The remaining participants reported using either a trackpad (*n* = 112), an optical mouse (*n* = 86), or a trackball (*n* = 14). A linear mixed model (LMM) did not show effects of Mouse Type on single-trial learning (STL), although we observed that participants using a trackpad exhibited longer reaction times than others, consistent with a previous report.(60)

Mouse position sampling rates depended on the exact hardware that each participant used to complete the task. Sampling rates were likely affected by features of the specific mouse used, along with features of the specific computer used, as computers may limit the rate at which the browser samples data in order to cope with limited processing power. In general, sampling rates were around 60 Hz (median ± interquartile range across all 213 online participants recruited: 62.46 ± 2.17 Hz) but ranged from 19.23 Hz to 249.69 Hz. Note that the vast majority of sampling rates were near 60 Hz: Only 5% of sampling rates were < 41.79 Hz, and only 5% of sampling rates were > 126.65 Hz.

### Baseline training phase

For in-lab participants, the robot moved the participant’s hand to a central starting location (depicted by a grey circle) at the middle of the display while hand and cursor feedback were hidden. They were instructed to hold their hand still in the starting location util the target turned green, at which point they should make a straight slicing movement through the target. After a 100 ms delay, the robot moved the hand back to the starting location. Participants completed 5 of these trials with online and endpoint cursor feedback, followed by 5 trials without visual feedback of the cursor location. Endpoint feedback was constituted by the cursor remaining at the position where it had passed the target radius for 50 ms. Participants then completed 10 alternating trials on which the target turned green and stayed green (Execution, ‘Go’ trials) and on which the target turned magenta 100 ms after turning green, signaling that participants should withhold their movement (No-Movement, ‘Stop’ trials). After this baseline phase, participants were instructed to continue following these instructions for the remainder of the experiment.

Online participants experienced an identical baseline phase, with the exception that they were instructed to move their mouse into a central starting location on the first trial and subsequently saw their mouse cursor reappear near the starting location 100 ms after the completion of the reaching movement, so that participants could quickly return to the start location to initiate the next trial.

### Test phase: Rotation and Error-Clamp Experiments (Fig. 2, Supplemental Fig. 2, Supplemental Fig. 3)

During the test phase, 480 (in-lab) or 270 (online) total trials were divided into 3-trial triplets (**Fig. 1C**). The first and last trials of all triplets were Go trials, and participants received neither online nor endpoint feedback about cursor location on these trials. The second trial of each triplet was either a Movement or a No-Movement trial. On Movement trials, participants either received rotated/error-clamped(22) visual feedback (15° clockwise [-, CW] or counterclockwise [+, CCW], with sign randomized across trials) or veridical/0° error-clamped visual feedback of their cursor location. On No-Movement (Stop) perturbation trials, participants viewed a brief animation of the cursor moving straight to the center of the target following a trajectory deflected by ±15° from the target center. Animation onset latency was set as a running median of the participant’s reaction times on the previous 5 trials, and animation duration was set as a running median of the participant’s movement times on the previous 5 trials. If a participant took longer than 400 ms to execute a movement, 800 ms to initiate the movement, their reach trajectory changed by >10° during the movement, or the reach terminated ≥ 60° away from the target, they received a warning and a 4s time-out. If a participant moved their hand (>5 mm in-lab [radius of the starting location]; anything >0 pixels online) on a No-Movement trial, the trial was immediately aborted, and they received a warning and a 4s time-out. The Stop manipulation was successful: Across the experiments, participants erroneously moved on 34.39 ± 20.63% (mean ± standard deviation) of Stop trials, suggesting that, for the most part, they were consistently planning movements on Stop trials.

For in-lab studies, we used 4 possible triplet perturbation trial types (Movement/No-Movement: ±15°), each of which occurred 40 times throughout each session. For online studies, we used 6 possible triplet perturbation trial types (Movement/No-Movement: ±15° or 0°), each of which occurred 15 times throughout each session. Triplets were pseudorandomly presented within each block, with the constraints that a single rotation (±15° or 0°) could not occur on more than 2 consecutive triplets and that the same movement condition (i.e., Movement or No-Movement) could not occur on more than 3 consecutive triplets. Three repetitions of each triplet type occurred in blocks of 18 triplets, and participants received a break after each of these blocks.

### Test phase: Rotation and Error-Clamp Experiments with 0°

*Perturbations on Movement Trials (Fig. 3)*. Experiments were conducted as described above for the other online experiments, with the exception of the details described in this section. For the experiments described in **Fig. 3A-D**, we used a reduced set of 3 possible triplet Perturbation trial types (No-Movement, 15° clockwise error; No-Movement, 15° counterclockwise error; Movement, 0° rotation). We maintained an equal number of Movement and No-Movement triplets throughout the session in order to ensure that participants would reliably respond to the “Go” cue presented at the start of each trial. So, each No-Movement triplet type occurred 22 times, while the Movement triplet type occurred 44 times. Triplets were pseudorandomly presented within each block, with the constraints that a single non-zero rotation (15° clockwise, 15° counterclockwise) could not occur on more than 2 consecutive triplets.

For the experiments described in **Fig. 3E-H**, we used a set of 4 possible triplet Perturbation trial types (No-Movement, 15° clockwise error; No-Movement, 15° counterclockwise error; No-Movement, 0° error; Movement, 0° error-clamp). To maintain an equal number of Movement and No-Movement triplets throughout the session, each No-Movement triplet type occurred 15 times and the Movement triplet type occurred 45 times.

### Questionnaire

As we could not receive verbal confirmation that participants understood the task instructions in the online version of the task, we asked subjects to fill out a brief questionnaire to query their understanding of the task. The questionnaire asked participants to attest whether or not 1) their goal was to move the real mouse and not the cursor straight through the green targets and whether or not 2) their goal was to move the white cursor (not the real mouse) straight through the green targets. Participants could select the options, “True,” “False,” or “Not Sure.” Participants were considered to have understood the instructions if they answered both questions correctly (i.e., answered “True” to question 1 and “False” to question 2). The majority of participants answered both questions correctly (138 of 213 participants [65%]), suggesting that most participants understood the task instructions. Nonetheless, these participants made up the dataset for all reported analyses for the online experiments, and all other online participants were excluded from analyses to exclude potential effects of explicit re-aiming.

### Data analysis

Data were processed in Python 3.8.5 and Matlab 2018a. Trials with movement were excluded from analysis 1) if any of the reaches in the triplet were not straight (aspect ratio > participant-wise mean + 3 * participant-wise standard deviation), 2) if the participant received any warning for failure to follow task instructions (see *Feedback for failure to follow task instructions*, above), or 3) if the triplet included a No-Movement No-Go perturbation trial with any detectable mouse movement (>0 pixels online, >5 mm in lab).

Reach endpoint angle was computed as the angular distance between the center of the target and the point at which the mouse passed the target’s radial distance. Because mouse sampling rates did not always allow us to measure mouse position at the exact target radius during the online study, we used the last sample before and the first sample after the mouse passed the target radius to compute an interpolated mouse position at the target radius, as described in a previous report.^48^ We note that analyses comparing these measures to measurements at the last sample of the reach (even when it was beyond the target) or the hand angle at peak velocity did not result in substantially different hand angle measurements or statistical outcomes.

Single-trial learning (STL) was measured as the difference between reach endpoint angle on the third and first trial of each triplet. For our initial analyses, the sign of STL corresponded to the direction of the relative change in hand angle, with clockwise changes in hand angle taking a negative sign and counterclockwise changes in hand angle taking a positive sign. When we collapsed STL data across rotation directions, we normalized the sign of STL so that changes in hand angle opposite the direction of the imposed rotation took a positive sign and changes in the direction of the rotation took a negative sign.

Remembered STL was quantified as the difference between reach endpoint angle on the first trial of one triplet and reach endpoint angle on the first trial of the previous triplet. When remembered STL is reported as a ratio, this value was computed by dividing remembered STL by the STL attributable to a given triplet.

### Statistics

Statistical tests were conducted in R (v. 4.0.3; packages rstatix(61), coin(62), MuMIn(63), lmerTest(64), lme4(65), r2glmm(66), emmeans(67), effsize(68), effectsize(69), magrittr(70), ggplot2(71), ggpubr(72), ggeffects(73)). The reproducible code and data are available at https://www.github.com/kimoli/LearningFromThePathNotTaken. Data from in-lab experiments were analyzed using a two-way repeated measures ANOVA. If an ANOVA showed a significant main effect or interaction, post-hoc pairwise tests were performed. When samples failed to satisfy the normality assumption of the pairwise t-test (assessed via a Shapiro-Wilk test), we used the more robust paired-samples Wilcoxon signed-rank test. Otherwise, we used the more powerful paired t-test. Effect sizes for ANOVA main effects/interactions were quantified via generalized 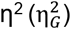, we quantified the effect sizes for t-tests using Cohen’s d, and we used the Wilcoxon effect size (r) to quantify effect sizes for signed-rank tests. For these and all subsequent analyses, we corrected for multiple comparisons using the false-discovery rate approach to maintain family-wise alpha at 0.05.

Data from the experiments conducted online did not satisfy multiple assumptions of the two-way repeated measures ANOVA (non-existence of extreme outliers, sphericity), so we employed a linear mixed modeling (LMM; R package lmerTest and lme4) approach for analysis of these data. All LMM’s included fixed effects of perturbation size and movement condition, as well as random effects of subject. Degrees of freedom were estimated using the Kenward-Rogers approach, and LMM outcomes were reported using ANOVA-style statistics. Partial R^2^ was computed to report effect sizes for the LMM factors (R package r2glmm). Post-hoc pairwise comparisons were performed between estimated marginal means computed from the LMM (R package emmeans).

For one-off comparisons between samples or to distributions with 0-mean, we checked samples for normality. When samples were normally distributed, we ran t-tests and computed Cohen’s d to report effect sizes for statistically significant results. Otherwise, we ran Wilcoxon-signed rank tests and measured effect sizes using the Wilcoxon effect size (r).

