## Supplementary Information for "Motor learning without movement"

### **Supplementary Information for**
Motor learning without movement.

Olivia A. Kim, Alexander D. Forrence, Samuel D. McDougle

Olivia A. Kim


**This PDF file includes:**

Figures S1 to S3
Tables S1 to S3
SI References

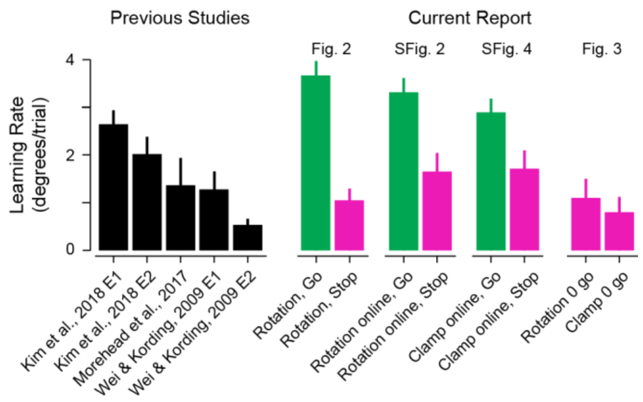


**Fig. S1. Learning rates reported in the literature and observed in the current study.** Learning rates for motor adaptation observed in previous studies are shown at left in black, and learning rates observed in each experiment in the current report are shown at the right, with data from Movement triplets shown in green and data from No-Movement triplets shown in magenta. Data are shown as mean ± SEM, and are shown for rotational/error clamp perturbations of 15°, with the exception of Wei & Kording, 2009 E2, where an 11° perturbation was applied. Papers referred to and their corresponding reference numbers: Kim et al., 2018 **(1)**; Morehead et al., 2017 **(2)**; Wei & Körding, 2009 **(3)**. “Rotation, Go” and “Rotation, Stop” show data from the in-lab experiment where participants saw 15° rotated feedback on Movement trials (i.e., data from Fig. 2), “Rotation online, Go” and “Rotation online, Stop” show data from the online experiment where participants saw 0-15° rotated feedback on Movement trials (i.e., data from Fig. S2). “Clamp online, Go” and “Clamp online, Stop” show data from the online experiment where participants saw 0-15° error-clamed feedback (i.e., data from Fig. S3). “Rotation 0 go” and “Clamp 0 go” show data from the online experiments where participants saw 0° perturbed feedback on Movement trials (i.e., data from Fig. 3). Abbreviations: E, experiment.

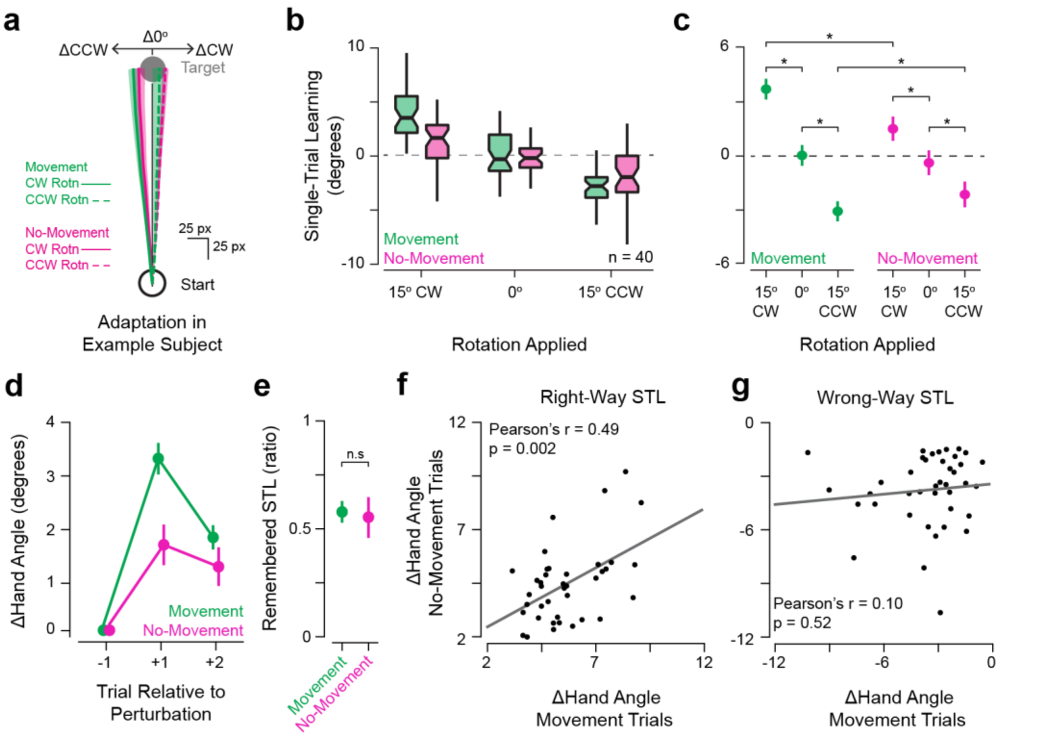


**Fig. S2. Single-trial learning in response to errors on Movement and No-Movement trials during an online visuomotor adaptation task.** (**a**) An example participant's mean ± SEM changes in reach paths across triplets (green: triplets with perturbations on Movement trials, magenta: triplets with perturbations on No-Movement trials, solid lines: perturbation was a CW rotation, dashed lines: perturbation was a CCW rotation). (**b**) Boxplot showing STL across Movement (green) and No-Movement (magenta) triplets for participants in an online version of the task described in Fig. 1 ($n$ = 40). (**c**) Estimated marginal means (EMMs) ± 95% confidence intervals from the linear mixed model (LMM) fit to participants' STL performance (summarized in **b**). The LMM (fixed effects: rotation [15° counterclockwise {CCW}, 0°, and 15° clockwise {CW}], movement condition [Movement, No-Movement], rotation x movement condition interaction; random effects: participant) revealed significant main effects of rotated cursor feedback ($F(2, 2223) = 136.46$, $p = 2.2 \times 10^{-16}$, *partial* $R^2 = 0.11$) and movement condition ($F(1, 2248) = 4.74$, $p = 0.03$, *partial* $R^2 = 0.002$), as well as a significant interaction ($F(2, 2229) = 12.40$, $p = 4.41 \times 10^{-6}$, *partial* $R^2 = 0.01$). Post-hoc pairwise comparisons of the EMMs from the model support the claim that rotated feedback induced a statistically significant degree of STL on both Movement (0° vs 15° CW: $t(2227) = 9.14$, $p_{adj} = 6.39 \times 10^{-19}$, *Cohen's* $d = 0.61$; 0° vs 15° CCW: $t(2220) = 7.81$, $p_{adj} = 2.61 \times 10^{-14}$, *Cohen's* $d = 0.52$) and No-Movement trials (0° vs 15° CW: $t(2225) = 3.92$, $p_{adj} = 1.39 \times 10^{-4}$, *Cohen's* $d = 0.31$; 0° vs 15° CCW: $t(2229) = 3.56$, $p_{adj} = 4.84 \times 10^{-4}$, *Cohen's* $d = 0.29$). Adaptation in the presence of a rotation was significantly greater in Movement trials than No-Movement trials for CW ($t(2238) = 4.98$, $p_{adj} = 1.26 \times 10^{-6}$, *Cohen's* $d = 0.37$) and CCW rotations ($t(2239) = 2.06$, $p_{adj} = 0.04$, *Cohen's* $d = 0.15$). Please refer to Table S1 for full details on post-hoc pairwise comparisons. (**d**) Group mean ± SEM change in (Δ) hand angle after exposure to Movement (green) and No-Movement (magenta) triplets' perturbations. Positive Δ values indicate that the change in hand angle proceeded opposite the direction of the perturbation (*i.e.,* in the direction that would counter the error). (**e**) Group mean ± SEM ratio of remembered STL to STL. Remembered STL was statistically significantly greater than 0 for both Movement (one-sample signed-rank test: $V = 819$, $p_{adj} = 1.09 \times 10^{-11}$, $r = 0.87$) and No-Movement triplets ($V = 769$, $p_{adj} = 9.69 \times 10^{-8}$, $r = 0.76$), but remembered STL did not significantly differ between movement conditions (paired-samples signed-rank test: $V = 441$, $p_{adj} = 0.68$). (**f**) Scatter plot showing the relationship between individual subjects' STL amplitude in the direction opposite the rotation on Movement and No-Movement trials. When we considered instances of STL in the direction that would compensate for the observed error (update opposite rotation, "Right-Way"), within-subject changes in hand angle were correlated between Movement and No-Movement trials (*Pearson's* $r = 0.49$, $p_{adj} = 0.002$). (**g**) As in **f**, but showing data from trials with changes in hand angle in the direction that would exacerbate the observed error (update in direction of rotation, "Wrong-Way"). These ΔHand Angle values were uncorrelated between Movement and No-Movement trials (*Pearson's* $r = 0.10$, $p_{adj} = 0.52$). These observations support the idea that the same learning process may underlie adaptive single-trial learning events in response to errors on both kinds of trials, while maladaptive changes in hand angle may be attributable to potential sources of random noise. Boxplot centers: median, notch: 95% confidence interval of the median, box edges: 1$^{st}$ and 3$^{rd}$ quartiles, whiskers: most extreme value within 1.5*interquartile range of the median. Statistical significance (* = $p < 0.05$; n.s. = $p \geq 0.05$) is. Abbreviations: STL – single-trial learning, Δ – change, CW – clockwise, CCW – counterclockwise.

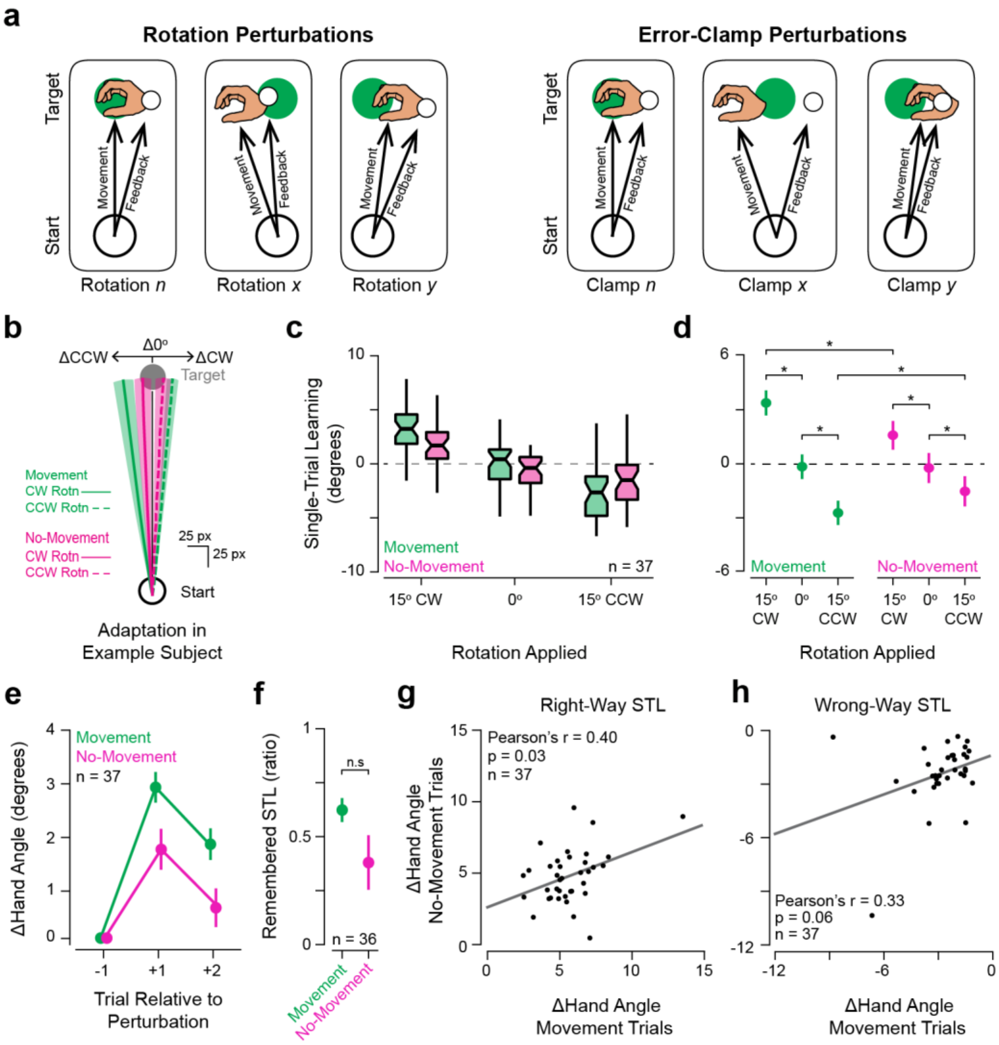


**Fig. S3. Single-trial learning in response to errors on Movement and No-Movement trials with error-clamped feedback or simulated errors.** (**a**) Diagrams showing the relationship between hand and cursor feedback movement directions under rotational (left) and error-clamp regimes (right). When rotations are applied, the cursor's movement direction is contingent upon the participant's movement direction. When error-clamp perturbations are applied, the cursor travels in a fixed direction, regardless of the direction that the hand travels. As error-clamp perturbations render deliberate changes in movement direction useless, they are often used in studies attempting to isolate implicit motor adaptation processes. (**b**) An example participant's mean ± SEM changes in reach paths across triplets (green: triplets with perturbations on Movement trials, magenta: triplets with perturbations on No-Movement trials, solid lines: perturbation was CW error-clamp, dashed lines: perturbation was CCW error-clamp). (**c**) Boxplot showing STL across Movement (green) and No-Movement (magenta) triplets for participants ($n$ = 37) in an online experiment where cursor feedback was error-clamped on Movement trials. (**d**) Estimated marginal means (EMMs) ± 95% confidence intervals from the linear mixed model (LMM) fit to participants' STL performance (summarized in **c**). The LMM (fixed effects: rotation [15° counterclockwise {CCW}, 0°, and 15° clockwise {CW}], movement condition [Movement, No-Movement], error-clamp x movement condition interaction; random effects: participant) revealed significant main effects of error-clamped cursor feedback ($F$(2, 1829) = 79.46, $p$ = 2.2 x $10^{-16}$, *partial* $R^2$ = 0.08) and an interaction between error-clamp and movement condition ($F(2, 1832) = 8.45$, $p = 0.0002$, *partial* $R^2 = 0.0003$), although there was no main effect of movement condition ($F(1, 1844) = 0.60$, $p = 0.44$). Post-hoc comparisons of the EMMs from the model revealed significant STL in response to non-zero error-clamped feedback on both Movement (0° vs 15° CW: $t(1827) = 7.55$, $p_{adj} = 3.08 \times 10^{-13}$, *Cohen's d* = 0.56; 0° vs 15° CCW: $t(1828) = 5.57$, $p_{adj} = 8.84 \times 10^{-8}$, *Cohen's d* = 0.41) and No-Movement trials (0° vs 15° CW: $t(1830) = 3.21$, $p_{adj} = 0.002$, *Cohen's d* = 0.29; 0° vs 15° CCW: $t(1832) = 2.25$, $p_{adj} = 0.03$, *Cohen's d* = 0.22). Adaptation in the presence of a 15° error-clamp was significantly greater on Movement trials than No-Movement trials for CW ($t(1846) = 3.49$, $p_{adj} = 0.0009$, *Cohen's d* = 0.29) and CCW clamps ($t(1846) = 2.29$, $p_{adj} = 0.03$, *Cohen's d* = 0.19). Please refer to Table S2 for further details on post-hoc comparisons in this panel. (**e**) Group mean ± SEM change in (Δ) hand angle one and two trials after exposure to Movement (green) and No-Movement (magenta) triplets' perturbations. Positive Δ values indicate that the change in hand angle proceeded opposite the direction of the perturbation (*i.e.,* in the direction that would counter the error). (**f**) Remembered STL shown as the ratio of relative hand angle 2 trials after experiencing a perturbation to the relative hand angle 1 trial after the perturbation (STL). Remembered STL was significantly greater than 0 after both Movement (green; one-sample t-test: $t(36) = 11.31$, $p_{adj} = 6.23 \times 10^{-13}$, *Cohen's d* = 1.86) and No-Movement triplets (magenta, one-sample signed-rank test: $V = 579$, $p_{adj} = 5.95 \times 10^{-5}$, $r = 0.64$), but did not exhibit statistically significant differences between movement conditions (paired t-test: $t(35) = 1.71$, $p_{adj} = 0.09$). Remembered STL on No-Movement trials could not be computed for one participant, so $n = 36$ instead of 37 in this panel. (**g**) Scatter plot showing the relationship between individual subjects' STL amplitude in the direction opposite the error-clamp on Movement and No-Movement trials (i.e., the "Right-Way"). Right-way changes in hand angle were correlated between Movement and No-Movement trials (*Pearson's* $r = 0.40$, $p_{adj} = 0.03$). (**h**) As in (**g**), but showing data from trials on which STL proceeded in the same direction as the error-clamp (*i.e.*, the "Wrong-Way"). Wrong-Way changes in hand angle were not statistically significantly correlated between Movement and No-Movement trials ($r = 0.33$, $p_{adj} = 0.06$). Boxplot centers: median, notch: 95% confidence interval of the median, box edges: 1st and 3rd quartiles, whiskers: most extreme value within 1.5*interquartile range of the median. Statistical significance (* = $p < 0.05$; n.s. = $p \geq 0.05$) is indicated for selected comparisons. Abbreviations: STL – single-trial learning, Δ – change, CW – clockwise, CCW – counterclockwise.

**Table S1.** Pairwise post-hoc comparisons between estimated marginal means in Fig. S2c.

| Group 1 | | Group 2 | | Est. Diff. | t | df | FDR-adjusted p | Cohen's d |
|---|---|---|---|---|---|---|---|---|
| Trial Type | Rotation | Trial Type | Rotation | | | | | |
| M | 0° | M | 15° CW | -3.67° | -9.14 | 2227 | * $6.39 \times 10^{-19}$ | -0.61 |
| M | 0° | M | 15° CCW | 3.09° | 7.81 | 2220 | * $2.60 \times 10^{-14}$ | 0.52 |
| M | 15° CW | M | 15° CCW | 6.76° | 17.08 | 2223 | * $1.31 \times 10^{-60}$ | 1.13 |
| No-M | 0° | No-M | 15° CW | -1.88° | -3.92 | 2225 | * 0.0001 | -0.31 |
| No-M | 0° | No-M | 15° CCW | 1.75° | 3.56 | 2229 | * 0.0005 | 0.29 |
| No-M | 15° CW | No-M | 15° CCW | 3.64° | 7.41 | 2231 | * $3.86 \times 10^{-13}$ | 0.61 |
| M | 0° | No-M | 0° | 0.41° | 0.93 | 2240 | 0.35 | -- |
| M | 15° CW | No-M | 15° CW | 2.20° | 4.98 | 2238 | * $1.25 \times 10^{-6}$ | 0.37 |
| M | 15° CCW | No-M | 15° CCW | -0.93° | -2.06 | 2239 | * 0.04 | -0.15 |

*Note.* Abbreviations: M – Movement; No-M – No-Movement; Est. Diff. – Estimated Differences in degrees. Note that degrees of freedom pertain to the inputs to the LMM and are estimated using the Kenward-Rogers approach. Statistically significant p-values are indicated with asterisks. * $p < 0.05$.

**Table S2.** Pairwise post-hoc comparisons between estimated marginal means in Fig. S3d

| Group 1 | | Group 2 | | Est. Diff. | t | df | FDR-adjusted p | Cohen's d |
|---|---|---|---|---|---|---|---|---|
| Trial Type | Error-Clamp | Trial Type | Error-Clamp | | | | | |
| M | 0° | M | 15° CW | 3.54° | -7.55 | 1827 | * $3.08 \times 10^{-13}$ | -0.56 |
| M | 0° | M | 15° CCW | 2.57° | 5.57 | 1828 | * $8.84 \times 10^{-8}$ | 0.41 |
| M | 15° CW | M | 15° CCW | 6.11° | 13.14 | 1828 | * $8.51 \times 10^{-37}$ | 0.97 |
| No-M | 0° | No-M | 15° CW | -1.81° | -3.21 | 1830 | * 0.002 | -0.29 |
| No-M | 0° | No-M | 15° CCW | 1.29° | 2.25 | 1832 | * 0.03 | 0.20 |
| No-M | 15° CW | No-M | 15° CCW | 3.10° | 5.46 | 1833 | * $1.24 \times 10^{-7}$ | 0.49 |
| M | 0° | No-M | 0° | 0.08° | 0.16 | 1846 | 0.87 | -- |
| M | 15° CW | No-M | 15° CW | 1.81° | 3.49 | 1846 | * 0.0009 | 0.29 |
| M | 15° CCW | No-M | 15° CCW | -1.19° | -2.28 | 1846 | * 0.03 | -0.19 |

*Note.* Abbreviations: M – Movement; No-M – No-Movement; Est. Diff. – Estimated Differences in degrees. Note that degrees of freedom pertain to the inputs to the model and are estimated using the Kenward-Rogers approach. Statistically significant p-values are indicated with asterisks. * $p < 0.05$.

**Table S3.** Pairwise post-hoc comparisons between estimated marginal means in Fig. 3

| Group 1 | | Group 2 | | Est. Diff. | t | df | FDR-adjusted p | Cohen's d |
|---|---|---|---|---|---|---|---|---|
| Trial Type | Perturbation | Trial Type | Perturbation | | | | | |
| 0° Rotation Applied on Movement Trials | | | | | | | | |
| No-M | 15° CCW | No-M | 15° CW | 2.45° | 4.80 | 557 | * $2.07 \times 10^{-6}$ | 0.40 |
| 0° Error-Clamped Feedback on Movement Trials | | | | | | | | |
| No-M | 15° CCW | No-M | 15° CW | 2.37° | 4.32 | 805 | * 0.0001 | 0.37 |
| No-M | 15° CW | No-M | 0° | 1.36° | 2.48 | 801 | * 0.020 | 0.21 |
| No-M | 15° CCW | No-M | 0° | 1.01° | 1.83 | 800 | 0.067 | -- |

*Note.* Abbreviations: No-M – No-Movement; Est. Diff. – Estimated Differences in degrees. Note that degrees of freedom pertain to the inputs to the model and are estimated using the Kenward-Rogers approach. Statistically significant p-values are indicated with asterisks. * $p < 0.05$.
